# Hormone signaling and immune programs define differential endocrine responsiveness in high-risk breast tissue

**DOI:** 10.64898/2026.03.02.709108

**Authors:** Nadine Goldhammer, Marin Bont, Shruti Warhadpande, Michael Choi, Jose Cedano, Heather Greenwood, Julia Ye, Christopher J. Schwartz, Michael Alvarado, Cheryl Ewing, Karen Goodwin, Rita A. Mukhtar, Jasmine Wong, Shoko E. Abe, Julia Chandler, Jordan Jackson, Olufunmilayo I. Olopade, Michael J. Campbell, Allison Lam, Chaelee Park, Anna Vertido, Laura J. van ‘t Veer, Nola Hylton, Laura J. Esserman, Jennifer M. Rosenbluth

**Affiliations:** Department of Medicine, University of California, San Francisco, USA; Department of Medicine, University Medical Center Utrecht, The Netherlands; Department of Surgery, University of California, San Francisco, USA; Department of Radiology and Biomedical Imaging, University of California, San Francisco, USA; Department of Pathology, University of California, San Francisco, USA; Memorial Sloan Kettering Cancer Center, New York, USA; Center for Clinical Cancer Genetics and Global Health, University of Chicago, USA; Department of Laboratory Medicine, University of California, San Francisco, USA; Chan Zuckerberg Biohub, San Francisco, USA

**Author notes:** These authors contributed equally (Nadine Goldhammer, Marin Bont).

## Abstract

Hormone therapies are frequently used to reduce breast cancer risk in individuals at increased risk for primary or subsequent disease; however, tissue-level responses to these therapies are heterogeneous and incompletely understood. Background parenchymal enhancement (BPE) on breast magnetic resonance imaging (MRI) provides a non-invasive radiologic readout of breast tissue features associated with endocrine responsiveness and cancer risk. Although BPE is associated with hormonal exposure, a subset of patients with BPE do not show a response to preventive endocrine therapy and therefore may remain at increased breast cancer risk. In this study, we integrated single-nucleus RNA sequencing and spatial transcriptomics to define the determinants of endocrine responsiveness in the setting of BPE. We identify hormone-driven epithelial cells with high levels of estrogen signaling and endocrine responsiveness, together with immune-associated epithelial programs characterized by diminished luminal identity and increased expression of immune-modulatory pathways, including major histocompatibility complex (MHC) class II and CD74. Functional organoid assays validate that these epithelial states exhibit differential sensitivity to tamoxifen and demonstrate that inflammatory signals can induce immune-modulatory epithelial programs. Together, our findings identify hormone signaling and immune programs as key determinants of endocrine responsiveness in breast tissue and provide a biological basis for interpreting radiologic markers relevant to cancer prevention.

## INTRODUCTION

Breast cancer is the most commonly diagnosed cancer in women, with a lifetime risk of one in eight.^1^ Although mammographic screening has improved early detection^2^, rates of advanced-stage disease remain largely unchanged.^3,4^ In addition, breast cancer recurs in 10-30% of patients within 10 years following treatment.^5–7^ To achieve meaningful reductions in incidence and mortality, personalized prevention strategies for primary and recurrent breast cancers guided by biologic indicators of risk are needed. Accordingly, we sought to evaluate radiologic biomarkers in individuals at elevated breast cancer risk, including cancer-naïve individuals with established risk factors as well as participants with prior ductal carcinoma *in situ* (DCIS) or invasive breast cancer, reflecting the ongoing risk of subsequent breast cancer in these populations.

Among proposed imaging biomarkers of breast cancer risk, mammographic density (MD) on mammography and background parenchymal enhancement (BPE) on magnetic resonance imaging (MRI) are two well-established independent predictors.^8–10^ MD refers to the amount of dense tissue on a mammogram and is classified into four major categories: A (almost entirely fatty), B (scattered areas of fibroglandular tissue), C (heterogeneously dense), and D (extremely dense). BPE categorizes the degree of fibroglandular tissue enhancement by contrast as low (minimal (<25%) or mild (25-50%)), or high (moderate (50-75%), or marked (>75%)).^11^ Both MD and BPE are modifiable with endocrine therapy^12–15^ and hold potential as biomarkers of therapeutic response in breast cancer prevention, including risk reduction for subsequent disease. Current clinical trials are investigating the utility of non-invasive imaging features for predicting treatment outcomes and informing personalized prevention strategies.^16–27^ However, the biologic foundations of these imaging features and how they reflect underlying cellular states associated with risk remain largely unknown.

Previous studies have shown that the breast tissue of women with high MD contains a higher proportion of fibroblasts and epithelial cells, with differences in extracellular matrix and active crosstalk between those cell types that may contribute to increased proliferation and tumorigenesis.^26,28–30^ The precise relationship of estrogen exposure to this process is a subject of debate.^31–34^ In contrast, the cellular and molecular mechanisms underlying BPE remain largely unexplored, despite its consistent association with breast cancer risk.

BPE is influenced by both endogenous factors (menstruation, menopausal status) and exogenous hormone exposures (endocrine therapy, hormone replacement therapies (HRT), and hormonal contraceptives).^12,35–40^ Yet, responses to estrogen-modulating therapies vary widely: not all women who take tamoxifen for risk reduction experience a reduction in BPE on MRI, and a lack of BPE response is thought to reflect reduced preventive efficacy.^12,13,41^ Although estrogen receptor (ER) target genes are expressed in both epithelial and stromal cell types, the cellular and molecular mechanisms through which estrogen influences BPE and determines its responsiveness to endocrine therapies remain unknown.^42^

Because the molecular determinants of endocrine responsiveness in the preventive setting remain poorly understood, we investigated the cellular and transcriptional programs underlying the radiologic marker BPE to inform strategies for cancer risk stratification and personalized prevention. Using single-nucleus RNA sequencing and spatial transcriptomics on histologically normal breast tissue from women at elevated risk for breast cancer, including women with genetic susceptibility and prior history of DCIS or invasive cancer, we generated a comprehensive atlas of cell states associated with BPE in the high-risk setting. We show that high levels of BPE are associated with increased estrogen signaling and alterations in the immune microenvironment, and we functionally validate these pathways in patient-derived organoids (PDOs). Together, our results support a model in which BPE is associated with either high estrogen or inflammation, processes which can be targeted in personalized prevention approaches.

## RESULTS

### Generation of a transcriptomic dataset annotated for imaging-based breast cancer risk factors

To gain a comprehensive understanding of cellular and molecular mechanisms associated with imaging-based breast cancer risk factors, we collected primary normal human breast tissues from women at elevated breast cancer risk, including individuals with genetic susceptibility, and/or a history of DCIS or invasive cancer, representing risk for both first and subsequent breast cancer events. Non-cancerous breast tissue specimens from women undergoing breast surgery were used for multi-level analyses encompassing transcriptomic, protein-level, and functional assessments (Fig. 1A). Single-nucleus RNA sequencing was performed on specimens annotated for BPE, MD, age, body mass index (BMI), menopausal status and use of HRT, and parity using the 10x Chromium workflow (Fig. 1B). After quality control, including regression of ambient and mitochondrial RNA, 41,376 nuclei from 31 women were analyzed.^43,44^

**Figure 1:**
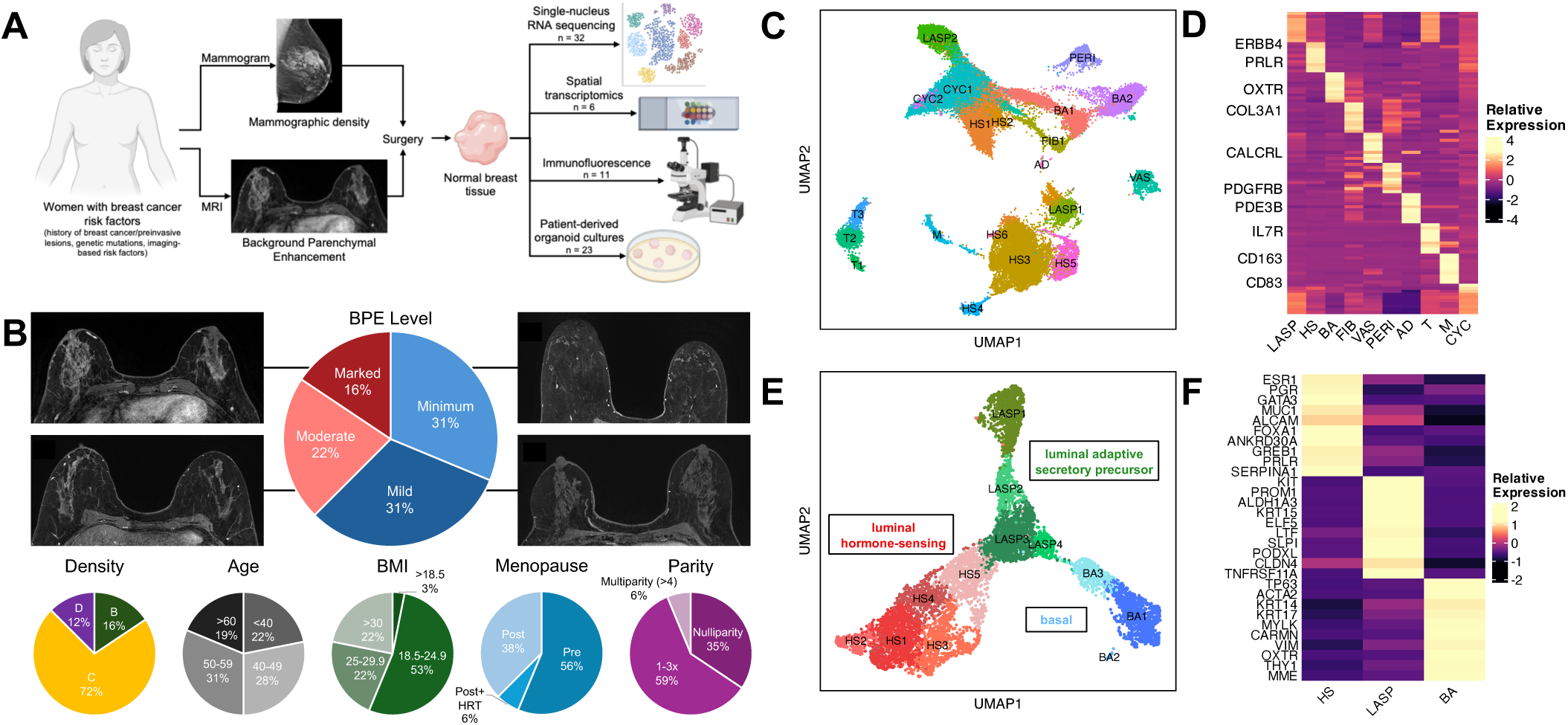
Generation of a breast snRNA-seq dataset annotated for imaging-based breast cancer risk factors. A) Cellular and molecular differences underlying imaging-based breast cancer risk factors were investigated at the transcriptomic, protein, and functional levels. Schematic shows overview of study workflow. B) Normal human breast tissues for single-nucleus RNA sequencing reflect patient heterogeneity. (Upper) Pie plot showing distributions of patients with distinct BPE levels and representative MRI images of the four BPE categories. (Lower) Pie plots showing the distribution of primary and secondary clinical annotations across the dataset. C) Single-nucleus RNA sequencing of primary breast specimens detects the main cell types of the normal human breast. UMAP plot showing nuclei from 31 women grouped into 20 clusters (color-coded) representing ten major cell types/states. AD = adipocytes, BA = basal cells, CYC = cycling cells, FIB = fibroblasts, HS = hormone-sensing cells, LASP = luminal adaptive secretory precursor cells, M = macrophages, PERI = pericytes, T = T cells, VAS = vascular cells. D) Expression of cluster-specific DEGs in normal human breast tissues. Heatmap showing the relative expression of the top ten highest expressed genes per cell type. E) Unsupervised clustering of epithelial cells from normal human breast tissues. UMAP plot showing 7,842 epithelial cells after re-clustering that reflect the three main epithelial cell types (HS = red, LASP = green, BA = blue) of the normal human breast. F) Expression of epithelial lineage marker genes in epithelial cells of normal human breast tissues. Heatmap showing the relative expression of selected epithelial breast lineage markers in the three main epithelial clusters.

Unsupervised clustering resulted in twenty clusters representing ten main cell types of the normal human breast (Fig. 1C-D, Suppl. Fig. 1A). The ten epithelial cell clusters each corresponded to one of the three main epithelial cell lineages of the normal breast^23–25^, namely hormone-sensing (HS) cells (also known as mature luminal cells), luminal adaptive secretory precursor (LASP) cells (also known as luminal alveolar or luminal progenitor cells), and basal (BA) cells (also known as myoepithelial cells) (Fig. 1C-D, Suppl. Fig. 1A).^48,49^ Stromal clusters represented T cells, macrophages, adipocytes, vascular cells, fibroblasts, and smooth muscle cells/pericytes (Suppl. Figure Fig. 1C-D1A).^30^

Each sample contained nuclei from various cell types in different ratios (Suppl. Fig. 1B), consistent with expected and widely reported patient-to-patient heterogeneity of cell types in the human breast.^45,46,48,51^ To further investigate stromal and epithelial cell populations, epithelial nuclei from the two groups were re-clustered and analyzed separately resolving transcriptionally distinct cell states, including hormone-sensing and immune cell subsets (Fig. 1E-F and Suppl. Fig. 1C-E).

Together, these analyses establish a comprehensive single-nuclei dataset from non-malignant human breast tissues of patients with genetic predisposition and/or a history of DCIS or invasive cancer, representing a cohort at elevated risk for primary and recurrent disease. Using this resource, we next analyzed transcriptomic changes associated with radiologic features of breast cancer risk.

### High mammographic density is associated with increased collagen expression

We first examined molecular changes associated with MD. However, because our high-risk cohort predominantly included samples with heterogeneous (C) MD, we lacked sufficient power for a formal comparative analysis. Nonetheless, our limited analysis was consistent with findings from previous studies: over-representation analysis of hallmark gene sets annotated in the Molecular Signatures Database (MSigDB)^52^ confirmed reports of increased tumor necrosis factor α (TNFα) signaling in pseudo-bulked HS and basal epithelial cells in the setting of high MD (Suppl. Fig. 2A).^53^ In addition, we assessed genes recently reported to be associated with high tissue stiffness and MD.^26^ We confirmed increased expression of extracellular matrix genes in samples with extreme compared to scattered MD (Suppl. Fig. 2B). Comparing epithelial cells from tissues with extreme MD to tissues with scattered MD we observed increased levels of genes encoding collagens (B = -0.022; D = 0.003), specifically members of the collagen VI family (B = -0.013; D = 0.013), and laminins (B = -0.135; D = 0.010; Suppl. Fig. 2C). Similarly, fibroblasts from tissues with extreme MD expressed significantly higher levels of collagens (B = 0.270; D = 0.433), especially fibrillar collagens (B = 0.397; D = 0.899), compared to scattered MD tissues (Suppl. Fig. 2D). We did not observe significant differences in cell type distribution across MDs (Suppl. Fig. 2E) Together, our limited analyses confirmed previous reports evaluating the molecular basis of MD.^26,54^ We next turned to the analysis of molecular mechanisms underlying background parenchymal enhancement (BPE), which remains largely unexplored.

### Estrogen-responsive programs define a subset of BPE-high tissues

To assess gene expression changes in tissues with distinct BPE levels, we compared cell type distribution and DEGs in from tissues with high (marked and moderate) versus low (mild and minimum) BPE levels using pseudo-bulk analysis (Fig. 2A-B). While we did not see any statistically significant differences in cell type abundance (Suppl. Fig. 2F), HS cells from tissues with high BPE levels were markedly enriched for estrogen signaling pathways (Fig. 2B-C). Tissues with high BPE levels had generally higher levels of estrogen response genes but not of ER itself compared to tissues with low BPE levels (Fig. 2C-D). We confirmed increased protein expression of the ER response gene *GREB1* by immunofluorescence protein staining in BPE-high (1.24 ± 0.19, n = 5) compared to BPE-low (0.80 ± 0.03, n = 6) tissues in a cohort matched for clinical annotations, including menopausal status (Fig. 2E-F). These differences in BPE high versus low tissues could be explained either by differential responsiveness of the tissue to estrogen, or by a difference in ER activity occurring in the tissue at the time the specimen was taken.

**Figure 2.**
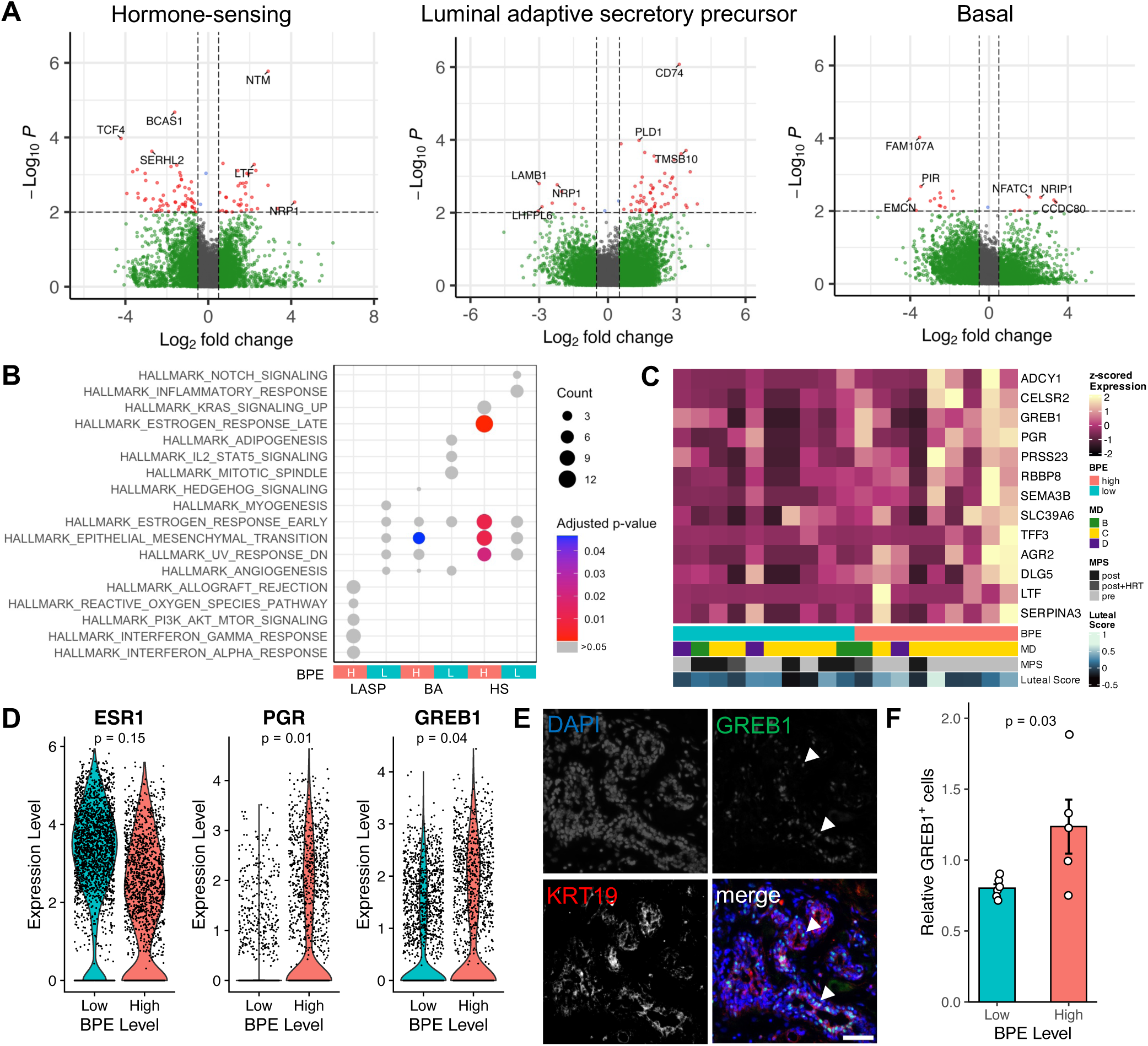
Estrogen signaling is increased in hormone-sensing cells from BPE-high tissues. A) Volcano plots showing differentially expressed genes between tissues with low (negative log2 fold change) and high (positive log2 fold change) BPE levels in HS (left), LASP (middle), and basal (right) epithelial cells. Each dot represents a gene and selected genes are labelled. Red = p value < 0.01 and log2 fold change > 0.5; blue = p value < 0.01 and log2 fold change < 0.5; green = p value > 0.01 and log2 fold change > 0.5; grey = p value > 0.01 and log2 fold change < 0.5. B) Hormone-sensing cells from tissue with high BPE levels are enriched for estrogen signaling pathways. Dot plot showing hallmark gene sets enriched for genes significantly upregulated in LASP, basal, and HS cells from tissue with high versus low BPE levels. C) Expression of genes associated with estrogen signaling in samples with distinct BPE levels. Heatmap showing expression of selected estrogen response genes (rows) in hormone-sensing cells across samples (columns). D) Expression levels of ER and estrogen response genes in tissues with distinct BPE levels. Violin plots showing expression of *ESR1*, *PGR*, and *GREB1* in tissues with low (blue) and high (red) BPE levels. P values based on DESeq2 test. E) GREB1 is expressed in luminal cells of the normal human breast. Representative fluorescence images showing breast tissue stained for nuclei (DAPI, blue), luminal keratin 19 (KRT19, red), and GREB1 (green). Arrows indicate GREB1+ nuclei. Scale bar = 50 μm. F) GREB1 protein expression in tissues with high compared to low BPE levels. Bar plot showing relative levels of GREB1+ cells in tissue sections from tissues with distinct BPE levels. Mean ± SEM; p value by t test.

### Breast organoids from BPE-high tissues show increased estrogen responsiveness in vitro

To functionally investigate whether estrogen-associated gene expression in tissues with high BPE levels is mediated by increased estrogen responsiveness, we generated PDOs from normal human breast tissues and optimized culture conditions to re-establish ER activity even after prolonged passaging. First, we compared two published organoid culturing media, type 1 and type 2 medium.^55–57^ Type 1 medium represents the original formulation for breast PDO culture, while type 2 is a modified version that contains additional growth factors such as β-estradiol, forskolin, and hydrocortisone. Organoids established and cultured in type 1 medium showed slower growth and morphologic differences compared to those cultured in type 2 medium (Suppl. Fig. 3A-B). Because of the faster growth rate in organoids cultured in type 2 medium, this culture condition was used for subsequent experiments. To ensure that all three major epithelial cell types are present in PDOs cultured in type 2 medium, single-cell RNA sequencing was performed on four PDOs after passaging seven to ten times. Unsupervised clustering of 33,201 cells resulted in 12 clusters representing HS, LASP, basal-luminal, basal, and cycling cells, confirming that the major epithelial cell populations were maintained with passaging (Fig. 3A-B).

**Figure 3.**
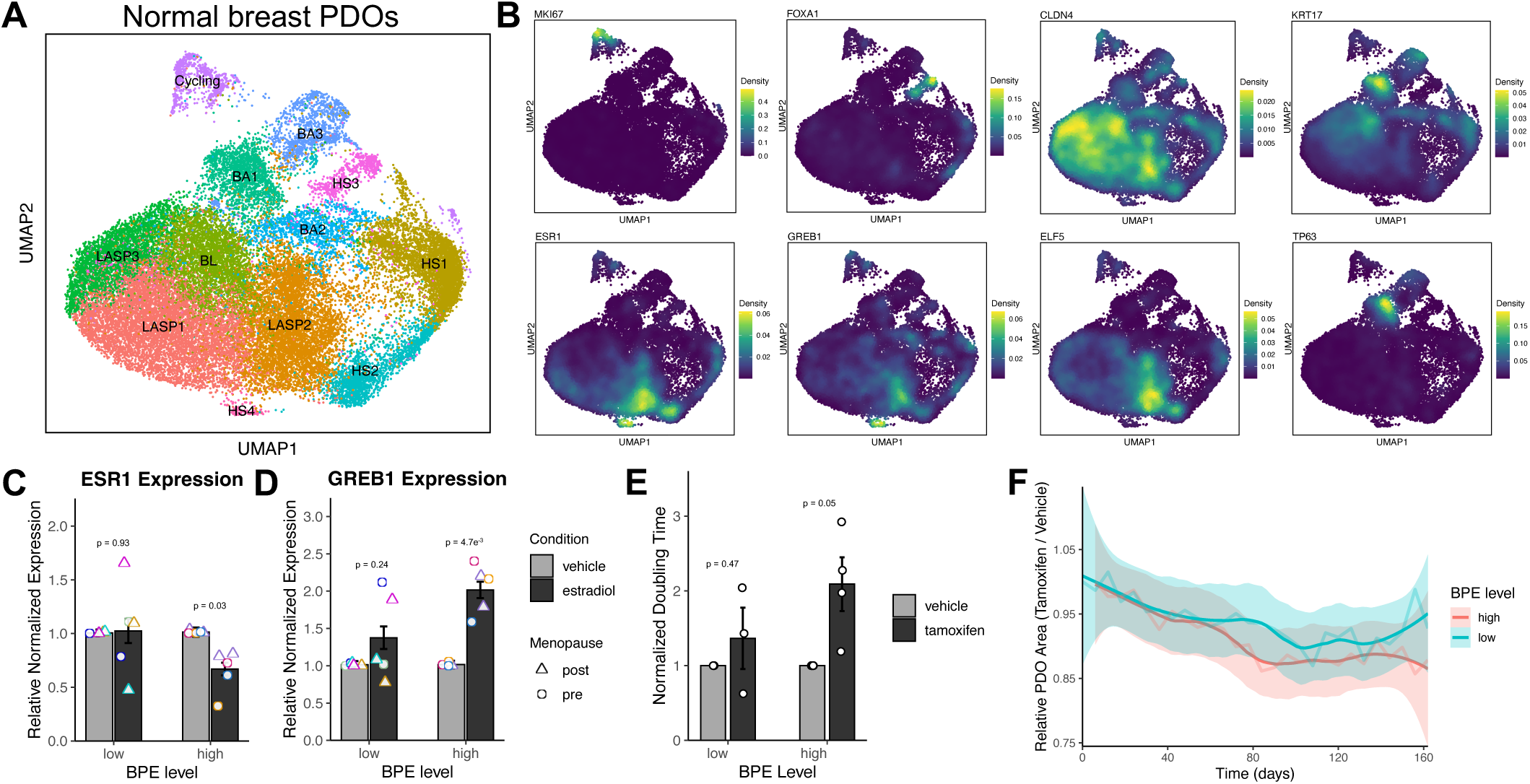
PDOs from women with high BPE levels have increased estrogen responsiveness *in vitro*. A) Clustering of cells from normal breast PDOs identifies different cell populations. UMAP plot showing 33,201 cells from four normal human breast PDOs assigned to 12 clusters representing epithelial cell types. B) Density plots showing expression of proliferation marker *MKI67*, hormone-sensing cell markers *ESR1*, *FOXA1*, and *GREB1*, LASP markers *CLDN4* and *ELF5*, and basal markers *KRT17* and *TP63*. C) Bar plot showing expression of estrogen receptor (*ESR1*) in PDOs from tissue with low or high BPE levels in response to stimulation with vehicle control or 100 nM β-estradiol. Each point represents an individual sample; colors indicate patient identity. Mean ± SEM, p values by t test with Benjamini-Hochberg correction, n = 5. D) Expression of estrogen response gene *GREB1* in PDOs from tissue with distinct BPE levels. Bar plot showing expression of estrogen response gene *GREB1* in PDOs from tissue with low or high BPE levels in response to stimulation with vehicle control or 100 nM β-estradiol. Each point represents an individual sample; colors indicate patient identity. Mean ± SEM, p values by t test with Benjamini-Hochberg correction, n = 5. E) Population doubling times in PDOs from tissues with high BPE levels in response to tamoxifen treatment. Bar plot showing relative population doubling times of PDOs from tissues with distinct BPE levels treated with vehicle control or 1 μM tamoxifen for seven days. Mean ± SEM, p values by t test. F) Tamoxifen treatment reduces PDO growth. Line plots showing area of PDOs from tissues with distinct BPE levels treated with 1 μM tamoxifen relative to vehicle control. Shown are averages of three PDOs with low BPE and four PDOs with high BPE.

Since type 2 organoid medium contains β-estradiol, leading to a downregulation of ER with time in culture, we next optimized a protocol to evaluate responsiveness to β-estradiol stimulation.^58–60^ The culture medium was depleted of combinations of β-estradiol, heregulin β1, EGF, and FGF-10, factors previously described to influence estrogen signaling.^61–63^ After seven days, PDOs were treated with 100 nM β-estradiol for four hours, and PDOs were harvested for RNA isolation to evaluate changes in the expression of estrogen response gene *GREB1* (Suppl. Fig 3C). *GREB1* has previously been described as an early estrogen response gene and is widely used to quantify estrogen activity *in vitro*.^64,65^ We found that depletion of multiple combinations enabled ER activity readout via an increase of estrogen-response gene *GREB1* in response to estradiol in an ER-dependent manner (Suppl. Fig. 3D and E). This increase was most reproducible across cultures and was statistically significant with the depletion of β-estradiol, heregulin β1, and FGF-10 (Suppl. Fig. 3D). All depletion conditions tested slowed PDO growth, however this occurred to a lesser extent when EGF was still present in the medium (Suppl. Fig. 3F). These findings established the conditions under which we could best assess estrogen responsiveness in PDOs, leading us to deplete β-estradiol, heregulin β1, and FGF-10 in subsequent experiments.

We used this protocol to compare estrogen responsiveness in five PDOs from patients with low BPE levels and from patients with high BPE levels. All PDOs were matched for age and menopausal status. Following stimulation, PDOs from high BPE but not low BPE tissues showed downregulation of ER expression in response to β-estradiol stimulation, likely due to an expected negative feedback regulation (Fig. 3C). In line with this, expression of the estrogen response gene *GREB1* was significantly upregulated only in PDOs from tissues with high BPE levels (Fig. 3D). These observations confirm increased responsiveness to estrogen in normal human breast tissues with high BPE levels.

Women at high risk of breast cancer commonly take tamoxifen for risk reduction.^66–68^ We investigated whether differences in estrogen responsiveness are associated with tamoxifen response. When treated with 1 µM tamoxifen or vehicle control, PDOs from tissues with high BPE levels showed a reduction in growth as shown by a significantly longer population doubling times (Fig. 3E), and reduced PDO area compared to vehicle control-treated PDOs (Fig. 3F). In addition, expression of *GREB1* was reduced in PDOs from high BPE tissues but not low BPE tissues in response to tamoxifen (Suppl. Fig. 3G). These results are consistent with reports that BPE is a predictive factor for breast cancer risk-reducing endocrine therapy.^21,23,24,69^

### Immune-associated epithelial programs characterize BPE-high tissue

We next evaluated gene expression in stromal cell populations of breast tissues with different BPE levels. Macrophages from BPE-high tissues upregulated genes related to inflammatory pathways, including genes associated with TNF⍺ signaling, IFNɣ response, and responsiveness to IL-2 and IL-6 cytokine networks, compared to BPE-low tissues, consistent with a chronically activated immune microenvironment (Suppl. Fig. 4A and Fig. 4A). In line with this, both vascular cells and LASP cells from tissues with high BPE levels showed an upregulation of interferon response pathways, though the differences in the latter did not reach statistical significance (Fig. 4A and 2B). In addition, since an inflammatory microenvironment can upregulate genes involved in antigen presentation, including aberrant MHCII expression, in normal or premalignant epithelium^70–72^, we compared the expression of MHC class I and II genes in tissues with varying BPE levels. Interestingly, we observed a significant upregulation of both MHCI and MHCII genes in BPE-high compared to BPE-low epithelial cells (Suppl. Fig. 4B-C).

**Figure 4.**
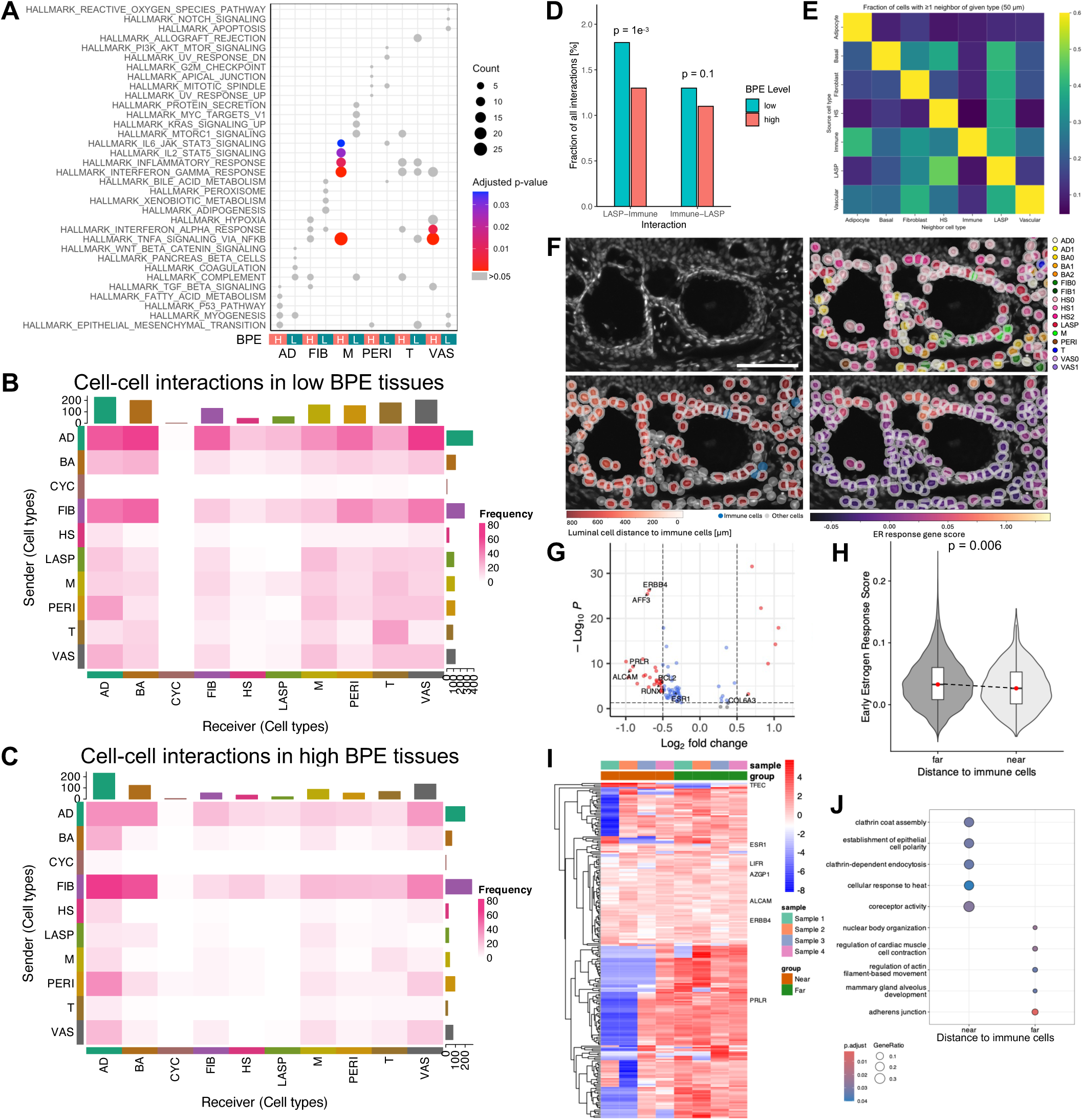
Inflammatory niches in BPE-high tissues alter epithelial-immune interactions and luminal identity. A) Macrophages from tissues with high BPE levels express genes associated with inflammatory pathways. Dot plot showing hallmark pathways enriched in stromal cell populations from BPE-low and BPE-high tissues. B) Cell-cell interactions in tissues with low BPE levels. Frequency plot showing interactions between the main cell types in the normal human breast in tissues with low BPE levels. C) Cell-cell interactions in tissues with high BPE levels. Frequency plot showing interactions between the main cell types in the normal human breast in tissues with high BPE levels. D) Interactions between LASP cells and immune cells in BPE-low and BPE-high tissues. Bar plot showing the percentage of total cell-cell interactions detected that involve LASP cells and immune cells (macrophages and T cells) across tissues with distinct BPE levels. P values by 2- sample test for equality of proportions with continuity correction. E) Spatial proximity between cell types in the normal breast. Heatmap showing the fraction of cells of each type that have at least one neighboring cell of the indicated type within a 50 μm radius. F) Representative fluorescence image of sample 4 (see Suppl. Fig. 4) stained for ssDNA and overlaid with cell segmentation annotating cell types, distance to immune cells, and ER response gene score by color. Scale bar = 100 µm. G) Gene expression differences between luminal epithelial cells close and far from immune cells in spatial transcriptomics data. Volcano plot showing adjusted p values and log2FC of DEGs between luminal cells within 50 μm (positive log2FC) versus distant (negative log2FC) from immune cells. H) Expression of early estrogen response genes in luminal cells in spatial relation to immune cells. Violin plot showing the module score of early estrogen response genes from the MSigDB in luminal cells distant (>50 μm) and proximal (<50 μm) from immune cells. Red dot indicates the median. p value by enrichment analysis with correction for multiple testing. I) Heatmap showing DEGs between luminal cells near to and far from immune cells, respectively, using a 50 μm cutoff. J) Dot plot showing enriched gene signatures in luminal cells near or far from immune cells.

To better understand the cellular implications of these changes, we next predicted cell-cell interactions in tissues from distinct BPE levels using the R package liana, which combines several cell-cell interaction methods to identify matching ligand-receptor cell-cell pairs.^73^ Although more nuclei from BPE-high samples were analyzed (30,540 nuclei from tissue with high levels of BPE versus 10,836 nuclei from tissues with low BPE levels), fewer cell-cell interactions overall were predicted in samples from BPE-high compared to BPE-low tissues (Fig. 4B-C). Investigating relative interactions between immune cells (macrophages and T cells) and the three main epithelial lineages in samples with different BPE levels, we observed that hormone-sensing and basal cells interact more with immune cells in tissues with high BPE levels compared to low BPE levels, as expected due to expression of high levels of genes involved in antigen presentation (Suppl. Fig. 4D). However, LASP cells interacted significantly less with immune cells in BPE-high versus BPE-low tissues (Fig. 4D). This was true despite similar levels of immune cell infiltrate (9.2% and 6.1% of total nuclei, respectively). Several studies suggest that LASP cells are the cell of origin for at least some breast cancer subtypes^45,74–76^ and reduced interactions with the immune system could allow pre-cancerous cells to evade immune detection and clearance.

To investigate whether LASP cells interact less with immune cells in BPE-high tissue because of changes in immune cell proximity to the epithelium, we performed spatial transcriptomics on sections of six normal breast tissues across patients with diverse oncology histories associated with high risk of breast cancer development or recurrence using the STEREO-seq platform which allows for whole transcriptome analyses at single-cell resolution (Suppl. Fig. 4E). Four samples passed quality control, and clustering was performed using the reference cell type expression profiles from our single-nucleus sequencing dataset, resulting in 16 distinct cell clusters (Suppl. Fig. 4E). The assessment of cellular neighbors of each cell showed that 7.6% of all LASP cells and 4.7% of all HS cells had at least one immune cell located within a 50 µm radius (Fig. 4E). In fact, multiple regions were notable for immune cells (T cells and macrophages) localized around terminal ductal lobular units (Suppl. Fig. 4E, Fig. 4F). This was confirmed when visualizing expression of estrogen signaling and inflammatory gene signatures (Suppl. Fig. 4E, Fig. 4F). Analysis of DEGs between luminal cells far (>50 µm) or near (<50 µm) to immune cells for each sample showed that luminal cells close to immune cells exhibit an altered cell state characterized by reduced expression of genes associated with luminal identity (Suppl. Fig. 4F, Fig. 4G-H).

Pseudo-bulk analysis confirmed that luminal cells proximal to immune cells expressed fewer genes associated with mammary identity including *ESR1*, *ALCAM*, and *ERBB4* (Fig. 4I-J). Few DEGs were detected in luminal cells proximal to immune cells (Fig. 4I). These included tumor suppressor genes such as *GAS7* and *TFEC*.^77,78^ This relative attenuation of luminal identity-associated transcription in immune-proximal cells suggests reduced engagement of estrogen-responsive programs.

### Inflammatory changes induce expression of immune-modulatory pathways in BPE-high tissues

Next, we further interrogated differences in gene expression between LASP cells in tissues with distinct BPE levels. Notably, CD74 was among the most significantly upregulated genes in LASP cells from high-BPE tissues compared with low-BPE tissues (Fig. 2A). CD74 regulates the binding and trafficking of major histocompatibility complex (MHC) class II molecules, and at high levels, such as those seen in inflammation or neoplasms, it is thought to not only inhibit the binding of endogenous peptides but also inhibit the binding and presentation of exogenous antigens.^79,80^ Furthermore, our analyses showed that LASP cells from BPE-high tissues, but not BPE-low tissues, interact with stromal cells by expressing CD74 (Suppl. Fig. 5A). CD74 expression in the normal breast epithelium was confirmed by immunofluorescence (Fig. 5A).

**Figure 5:**
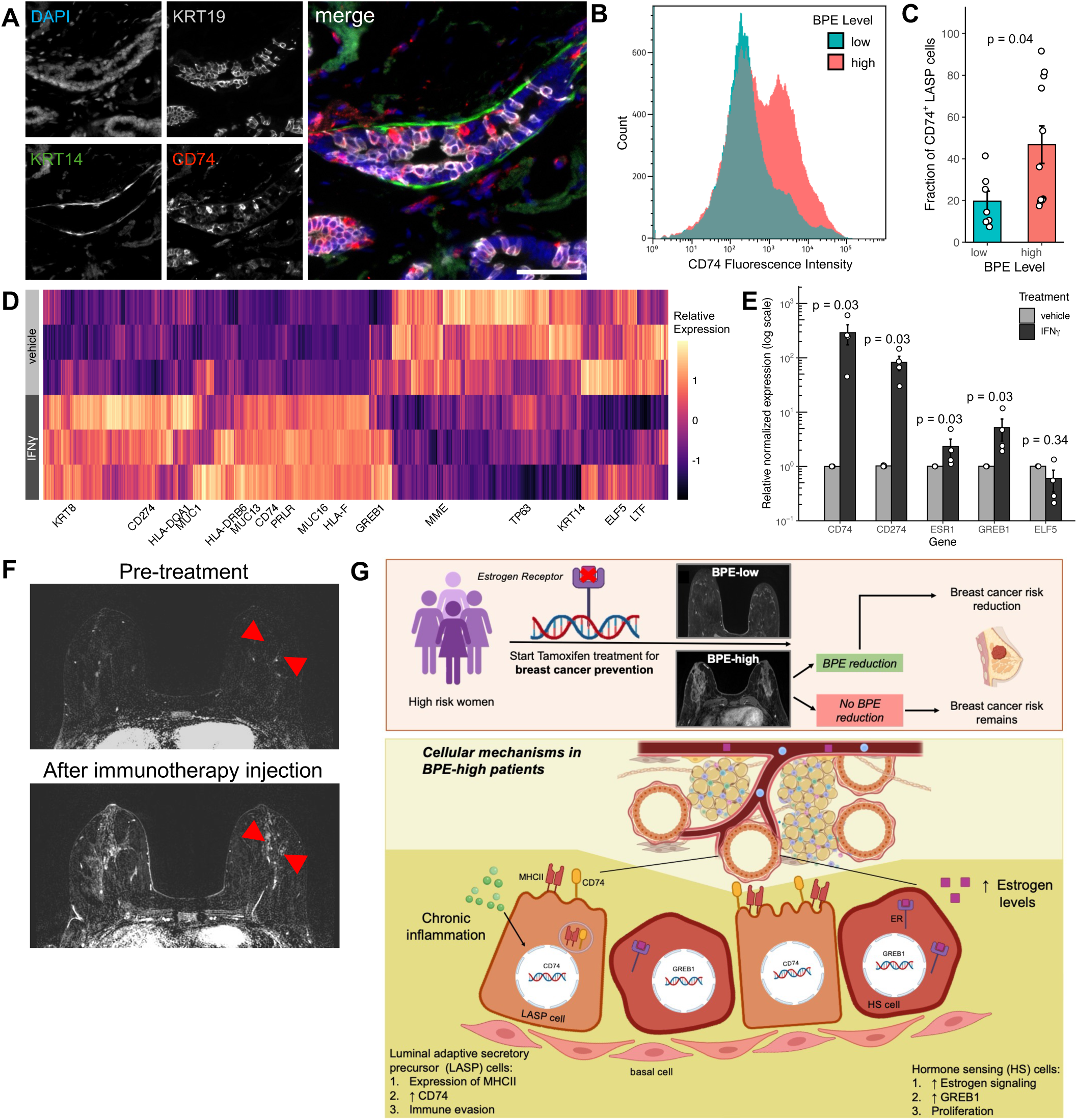
Inflammation induces immuno-modulatory cell states and is associated with increased BPE. A) CD74 is expressed in luminal cells of the normal human breast. Representative immunofluorescence images showing a normal human breast epithelial duct, where nuclei have been labelled with 4’,6-diamidino-2-phenylindole dihydrochloride (DAPI), myoepithelial cells are stained with an antibody against keratin 14, luminal cells are labelled with an antibody against keratin 19, and cells expressing CD74 are labelled using an anti-CD74 antibody. Scale bar = 50 μm. B) CD74 protein expression in LASP cells of PDOs from tissues with distinct BPE levels. Histogram showing fluorescence intensity of LASP cells from PDOs with distinct BPE levels stained for CD74. C) PDOs from tissues with high BPE levels are enriched for CD74^+^ LASP cells compared to PDOs from BPE-low tissues. Bar plot showing the fraction of CD74+ LASP cells in PDOs from tissues with distinct BPE levels. P value by Wilcoxon rank sum test. D) Differentially expressed genes between PDOs treated with IFNγ. Heatmap showing differentially expressed genes in PDOs treated with vehicle control (top, light gray) or 5 ng/ml IFNγ (bottom, dark grey). E) Expression of selected genes in response to IFNγ stimulation in PDOs. Bar plot showing expression of *CD74*, *CD274*, *ESR1*, *GREB1*, and *ELF5* in PDOs treated with vehicle control or 5 ng/ml IFNγ. P value by Wilcoxon rank sum test with Benjamini Hochberg correction. F) Representative post-contrast subtracted MR images showing a patient with mild BPE before treatment (top), whose BPE in both the left and right breasts increased to moderate after experimental injection of pembrolizumab and cytokines into the left breast (bottom). Red arrowheads indicate areas of BPE in the contralateral breast. G) Schematic overview showing proposed epithelial and immune cell changes in BPE-high tissues that may contribute to increased breast cancer risk.

To assess the functional role of CD74 in breast epithelial cells, we used PDOs derived from breast tissues with distinct BPE levels. Flow cytometric analysis showed that, in line with the results from single-nucleus RNA sequencing, CD74 protein expression was increased in LASP cells in an independent set of PDOs from tissues with high BPE levels compared to PDOs from tissues with low BPE levels (Fig. 5B-C). To assess if upregulation of CD74 could be a result of a pro-inflammatory environment in tissues with high BPE levels, we treated BPE-high PDOs with 5 ng/ml IFNɣ or vehicle control for 7 days (Suppl. Fig. 5B), resulting in a strong upregulation of *CD74* as well as *CD274*, encoding the immune evasion marker PD-L1 (Fig. 5D-E).^81,82^ We further observed that proteins of the MHC class II family, as well as other inflammatory pathways, were upregulated by IFNɣ in BPE-high PDOs (Fig. 5D and Suppl. Fig. 5C-D).

Interestingly, IFNɣ treatment also resulted in increased levels of *ESR1* and ER target genes such as *GREB1*, suggesting cross-talk between these pathways in this context^83–86^, while concomitantly reducing expression of other basal and secretory-alveolar LASP differentiation factors such as *ELF5* and *LTF* (Fig. 5D-E). Together, these results suggest that inflammatory signals can reprogram luminal cells to upregulate genes associated with immune evasion, such as CD74 and PD-L1, and downregulate differentiation factors, providing a potential mechanistic explanation for persistent enhancement despite endocrine therapy.

Further anecdotal evidence linking inflammation to BPE was identified via additional analysis of a tissue case from our recently reported phase I clinical trial evaluating intratumoral administration of the PD-L1 inhibitor pembrolizumab in combination with mRNA-2752 in patients with high-risk DCIS.^27^ The mRNA-2752 cocktail comprises a combination of IL-23, IL-36γ, and OX40L mRNAs, aiming to attract immune cells to the injection side. We observed an increase in BPE in the contralateral breast tissue after injection (Fig. 5F), which persisted for at least 7 months on follow-up MRIs obtained during both the luteal and follicular phases of the menstrual cycle. In combination with our transcriptomic and organoid-based studies, this result supports a model in which inflammation can induce BPE and increase breast cancer risk.

## DISCUSSION

Endocrine therapies can prevent primary and secondary breast cancer in a subset of patients. However, factors that determine which patients will benefit from preventive endocrine therapies and treatment options for patients who do not respond are currently lacking. Here, we characterized the breast cancer risk factor BPE, which has been demonstrated to be a potential surrogate marker for endocrine therapy responsiveness and risk reduction. We show that two mechanisms are mediating BPE on the molecular level: 1) increased estrogen responsiveness of HS cells and 2) upregulation of immune-modulatory pathways, including CD74 and PD-L1 in LASP cells as a result of chronic inflammation (Fig. 5G).

Preventive endocrine therapies are currently the only pharmaceutical interventions available for women at high risk for developing breast cancer and are similarly used long-term in patients with ER^+^ breast cancer to prevent recurrence. To develop new cancer prevention strategies, we need surrogate markers that correlate with risk reduction and can be used in clinical trials. BPE has long been recognized as an independent breast cancer risk factor, yet its underlying molecular and cellular determinants have remained largely undefined. Here, we identify distinct epithelial and immune-associated mechanisms underlying BPE, providing biological insight into this radiologic marker of risk, and potentially explaining why it is an independent risk factor from MD, which is associated with different biology, including changes in collagen and ECM.^26,87–91^

Previous observational studies demonstrate that hormone-associated states, including menstrual timing and menopause, are associated with high BPE levels.^12,24,35,36^ Using single-nucleus transcriptome sequencing, we confirmed increased estrogen signaling specifically in HS cells from BPE-high tissues and validated increased *GREB1* protein expression in an independent cohort of premenopausal patients. Notably, PDOs derived from BPE-high tissues retained heightened estrogen responsiveness and sensitivity to tamoxifen even after prolonged time in culture. Together, these findings indicate that BPE reflects intrinsic estrogen responsiveness of normal breast epithelium and suggest that estrogen-response signatures could be developed as predictive biomarkers in normal tissue, analogous to approaches used in invasive breast cancers.^92–97^

Not all patients with high BPE experience a reduction in enhancement in response to endocrine therapy, and inter-patient variability suggests that additional factors can influence BPE. Pro-inflammatory signatures in macrophages in BPE-high tissues were accompanied by expression of antigen presentation genes in LASP cells, including most notably the MHCII chaperone and immune modulator CD74. IFNɣ treatment induced MHCII genes, CD74, and PD-L1 in BPE-high PDOs, supporting a model in which epithelial cells adopt immune-evasive programs in response to a pro-inflammatory microenvironment. These immune niches could be detected by spatial transcriptomics across a small number of tissues representing diverse clinical contexts, and adjacent epithelial cells exhibited distinct changes in cell state reflective of decreased luminal differentiation pathways. Moreover, inflammatory processes can increase tissue vascularity and permeability^49^, consistent with inflammatory signatures in BPE-high vascular cells in our dataset, which may lead to contrast leakage and explain increased enhancement on MRI.

Together, our results point towards an inflammatory immune environment in tissues with high levels of BPE, a feature itself associated with breast cancer risk.^98^ Clinical trials of antiinflammatory agents such as low-dose aspirin have yielded mixed results^99,100^, underscoring the importance of appropriate patient selection. In this context, our findings suggest that anti-inflammatory therapy may be particularly beneficial for patients with persistent BPE despite tamoxifen therapy.^12^ In addition, several therapies targeting CD74 are currently under development for cancer treatment, and our data suggest that CD74 represents a candidate pathway for mechanistic investigation in preclinical models of breast cancer risk and prevention.^101,102^

Based on these findings, we propose a model in which high-risk patients with elevated BPE take tamoxifen for three to six months for risk reduction, after which their BPE response is assessed. A decrease in BPE suggests that the elevated signal was primarily driven by estrogen-responsive tissue changes, and these patients should remain on tamoxifen. In contrast, if BPE does not decrease, the elevation may instead reflect chronic inflammation, and patients may need to consider alternative approaches, such as anti-inflammatory therapies. This approach allows early assessment of therapeutic response, potentially sparing non-responders from the side effects of extended endocrine therapy and providing a more personalized risk-reduction strategy. These findings should inform the development of new response adaptive trial designs to optimize prevention in women at high risk.

## METHODS

### Sample and clinical data collection

Normal breast tissue samples from women undergoing breast surgeries for breast cancer, preinvasive lesions, or for preventive risk reduction were collected from the Breast Care Center at UCSF under IRB-approved protocols. Collected data included age, body mass index (BMI), breast cancer type if applicable, menopausal status, use of hormone replacement treatment (HRT; defined as estrogen containing HRT used within the last five years), treatment history, and imaging metrics. Specimens were collected either as single core biopsies or as pieces of surgically resected tissue, and visual tissue appearance ranged from fatty to dense. Upon arrival at the laboratory, a piece of each tissue was fixed in 10% formalin for the generation of formalin-fixed paraffin embedded (FFPE) tissue blocks for processing and review by a breast pathologist to confirm the absence of tumor histology. The residual normal human breast tissue chunks or core biopsies were minced into small pieces (1×1 mm), using a scalpel. The minced tissue was viably frozen in fetal bovine serum (FBS, HyClone, Cat. No. 89133-098) with 10% dimethyl sulfoxide (DMSO, Thomas Scientific, Cat. No. C752A55) and stored in liquid nitrogen as described previously or directly processed for PDO generation as described below.^55^

### Single-nucleus RNA sequencing, data processing, and analysis

snRNA-seq was performed on 32 normal breast tissues from women at high-risk for primary or recurrent breast cancer (due to genetic predisposition or history of breast lesions) using the 10x Chromium Nuclei Isolation Kit (Cat. No. PN-1000493) and the GEM-X Single Cell 3’ Kit v4 (Cat. No. PN-1000691) according to the manufacturer’s instructions. For sample preparation, we aimed to include both fatty and dense tissue regions whenever possible, up to a total of 85 mg. Tissues were lysed, debris removed, and nuclei were washed following the manufacturer’s instructions. The RNase inhibitor (NEB, Cat. No. M0314) was ordered separately.^46^ Single-nuclei suspensions were diluted in wash-and-resuspension buffer to concentrations ranging from 187-3750 nuclei/µl. Libraries were generated according to the manufacturer’s instructions. Libraries were sequenced on the Illumina NovaSeqX platform pooling libraries from eight samples and using four lanes with 5 billion clusters per library pool. Resulting FASTQ files were uploaded to the 10x Genomics cloud platform for alignment, filtering, barcode counting, and UMI counting using the Cell Ranger Count v8.0.1 pipeline.

Cell-free mRNA contamination was estimated to be 20% and removed from filtered count matrices using the R package SOUPx (v1.6.0).^44^ Resulting count matrices were analyzed using the R package Seurat (v5.1.0).^43^ Debris and duplets were removed by filtering nuclei with <200 and >6,000 genes and mitochondrial genes were regressed from the dataset as ambient RNA. Data was processed using the standard Seurat workflow. Briefly, data was log-normalized, variable features identified, data was scaled, and principal components analysis (PCA) was performed. Data from different samples were integrated using Harmony batch correction, followed by clustering and dimensional reduction.^103^ DEGs between clusters were identified using a Wilcoxon Rank Sum test. Clusters were annotated based on expression of DEGs and by label transfer using a reference dataset.^46^ All clusters except one that expressed adipocyte and epithelial genes could unambiguously be matched to previously described cell types. This cluster was included in the analysis of stromal and epithelial cells individually and contaminating cells were filtered during downstream analysis. Epithelial and stromal cells were subset from the dataset and analyzed separately following the same workflow. Due to the varying number of epithelial cells between samples, datasets were downsized to 10,000 nuclei per sample for analysis of epithelial cells. Pseudobulking to compare samples with distinct BPE levels was performed using Seurat’s AggregateExpression() function, and DEGs between BPE levels were determined based on a model using DESeq2, which uses a negative binomial distribution.^104^ Scores for the expression of MHC genes were calculated based on the following genes for MHC class I: HLA-A, HLA-B, HLA-C, HLA-D, HLA-E, HLA-F, and MHC class II: CD74, HLA-DRA, HLA-DPA1, HLA-DRB1, HLA-DPB1, HLA-DQB1, HLA-DQA1, HLA-DMA, HLA-DRB5, HLA-DMD, HLA-DQA2. The luteal gene signature was derived from Pardo et al.^105^ Enrichment analysis was performed using clusterProfiler (v4.12.6) with data from the MSigDB.^106,52^ Cell-cell interactions were analyzed using liana (v0.1.14).^73^ In addition, the following were used for data visualization: ggplot2 (v3.5.1), Nebulosa (v1.14.0), dittoSeq (v1.16.0), EnhancedVolcano (v1.22.0), and pheatmap (v1.0.12).^107–111^

### Generation and maintenance of PDOs

PDOs were generated as described previously.^55,56^ Briefly, fresh or freshly thawed minced normal human breast tissues were digested for 2-8 h at 37 °C with collagenase type 1 (ThermoFisher, Cat. No. 17101015) diluted in base medium. During the incubation period, the tissue was visually inspected and manually digested by pipetting with a 1 mL pipette every 30 minutes. Digestion was stopped by adding 2% FBS, and cells were spun down for 3 min at 300 x g. The cell pellet was resuspended in 50 µL Cultrex Basement Membrane Extract (BME) type 2 (BioTechne, Cat. No. 3532-010-02) and plated in a preheated (37 °C) 24-well non-cell culture treated plate (Corning, Cat. No. 3738). After 20 minutes, preheated (37°C) type II organoid expansion medium was added per well. Two expansion medium types were used as previously published, type 1 and type 2.^55,57^ PDO cultures were kept at 37 °C and 5% CO_2_. The medium was changed every 2-3 days. Organoids were passaged approximately every 1-4 weeks, according to their growth rate. For passaging, the medium was removed and PDO domes were gently resuspended in preheated (37 °C) TrypLE Express (Invitrogen, Cat. No. 12605036). After incubating at 37 °C for 2 minutes, digestion was stopped by transferring the PDO solution to ice-cold base medium. Note that PDOs were never digested to single cells to maintain the correct cellular architecture. PDOs were spun down for 3 min at 300 x g, and organoids were resuspended in BME and seeded as described above.

### Single-cell RNA sequencing

Single-cell RNA sequencing was performed on four PDO cultures grown in type 2 medium. Libraries were prepared using the Chromium 10x Genomics Single Cell 3’ Expression kit (v3) (Cat. No. PN-1000690) following the manufacturer’s instructions. Libraries were pooled and sequenced on the Illumina NovaSeq S4 platform. Alignment, filtering, barcode counting, and UMI counting was performed using the CellRanger pipeline on the 10x Genomics Cloud (v8.0.1). Filtered count matrices were analyzed using Seurat (v5.1.0). Debris and duplets were removed by filtering cells with <200 and >10,000 genes, and dying cells were removed by excluding cells expressing >25% mitochondrial genes. Data were then log-normalized, scaled, and principal components analysis (PCA) was performed. Data from different samples were integrated using canonical correlation analysis (CCA) integration^112^, followed by clustering and dimensional reduction. DEGs between clusters were identified using a Wilcoxon Rank Sum test. Density plots to visualize expression of certain genes were generated using Nebulosa (v1.14.0).

### Functional PDOs assays

To evaluate response to β-estradiol stimulation, PDOs were seeded in 96-well plates (Corning, Cat. No. 3595) and grown in type 2 medium until 70-80% confluency. Then, organoids were deprived of β-estradiol, heregulin β1, and FGF-10, and/or EGF for seven days. PDO growth was monitored using a Sartorius IncuCyte S3 live cell imaging system with the spheroid module. Growth factor depletion was followed by a four-hour treatment with 100 nM β-estradiol or ethanol as vehicle control. As a negative control, one organoid was additionally treated with the selective ER degrader (SERD) fulvestrant (Selleck Chemicals, Cat. No. 1191) to confirm ER dependence. After β-estradiol stimulation, PDOs were harvested in TRIzol (Fisher Scientific, Cat. No. 15596018) for RNA isolation. To assess the effects of treatment with interferon-ɣ or tamoxifen, PDOs were seeded in 96-well plates and treated for one week with 5 ng/ml interferon-ɣ (Gibco, Cat. No. AF-300-02), 1 µM tamoxifen (Cayman Chemical, Cat. No. 14854), or vehicle control. PDO growth was monitored using a Sartorius IncuCyte S3 live cell imaging system. After seven days, PDOs were harvested in TRIzol for RNA isolation. Harvested PDOs were either frozen in -80 °C or directly subjected to RNA isolation. Except where indicated, all experiments were performed in technical triplicate and in at least three PDOs from three different tissue donors.

### RNA isolation, reverse transcription, quantitative real-time PCR, and RNA sequencing

For RNA isolation, chloroform was added to PDOs previously harvested in TRIzol in a 1:5 ratio, and reaction mixtures were incubated at room temperature for 5 minutes. Phase separation was achieved by centrifugation at 12,000 x g for 15 minutes at 4°C. The aqueous phase containing RNA was harvested, and isopropanol was added in a 1:1 ratio. Reactions were incubated for ten minutes at room temperature and centrifuged at 12,000 x g for 10 minutes at 4°C. The supernatant was removed, and the pellet was washed with 100% ethanol, then centrifuged at 7,500 x g for 10 minutes at 4 °C to precipitate the RNA. The supernatant was removed, and the RNA pellet was dried for 5-10 minutes. The RNA was resuspended in DNase/RNase-free distilled water (Thermo Fisher, Cat. No. 10977015) and quantified using a Thermo Scientific NanoDrop microvolume spectrophotometer. RNA was stored at -80 °C.

The RNA-to-cDNA kit (Thermo Fisher, Cat. No. 4387406) was used for reverse transcription to generate cDNA following the manufacturer’s instructions. Quantitative real time PCR (qRT-PCR) was performed using the TaqMan Real Time PCR System. The following predesigned primer-probe sets were used (all Thermo Fisher Scientific): *ACTB* (Hs01060665_h1), *GAPDH* (Hs02786624_g1), *ESR1* (Hs01046816_m1), *GREB1* (Hs00536409_m1), *CD74* (Hs00269961_m1), *CD274* (Hs00204257_m1), and *ELF5* (Hs01063023_g1). Taqman Gene Expression Master Mix (Thermo Fisher, Cat. No. 4369016) with primers and 10 ng cDNA were combined in a 96- or 384-well PCR reaction plate (Applied Biosystems, Cat. No. N8010560 and 4309849). PCR reactions and quantitative measurements were performed using the QuantStudio™ 6 Flex System. Data were exported to Excel, and all relative gene expressions were determined using the comparative cycle threshold (CT) method. Statistical analyses were performed in R (v4.5.1). Gene expression differences between conditions were assessed by t-tests, and p-values were adjusted for multiple testing using the Benjamini-Hochberg procedure if applicable.

RNA sequencing was performed by Novogene on a NovaSeq X Plus system. Data quality control, filtering, and alignment to a reference genome were performed by Novogene. Raw gene-level count matrices were filtered to remove lowly expressed genes, and differential expression analysis was performed using DESeq2 (v1.48.2). Variance-stabilized counts were used for heatmap visualization, and significantly DEGs (padj < 0.05, |log₂FC| > 1) were used for gene set enrichment analysis with clusterProfiler (v4.16.0).

### Immunofluorescence staining

Tissue pieces were fixed in 10% formalin (Electron Microscopy Sciences, Cat. No. 15743-120) overnight and stored in 70% ethanol until embedding. Fixed tissues were embedded in paraffin following standard protocols. Sections of 8-10 µm thickness were cut using a microtome and sections were dried for 1 hour at 60 °C using a Techne Hybridiser HB-1D. Sections were deparaffinized by two incubations with xylene (RPI, Cat. No. 111056-CS) for 5 min, followed by rehydration using an ethanol series. Antigen retrieval was performed by incubating sections for 40 min in 1x citrate buffer (pH = 6, abcam, Cat. No. ab93678) in a steamer. Slides were allowed to cool to room temperature for 20 min and washed once with water. Tissues were permeabilized by incubation with 0.5% Triton-X100 (Sigma Aldrich, Cat. No. X100) for 10 min at 4 °C. After two washes, tissues were blocked using 10% goat serum (Gibco, Cat. No. 16210-064) in tris-buffered saline with tween (TBS-T, Apex, Cat. No. 18-235) with 1% bovine serum albumin (BSA, Sigma Aldrich, Cat. No. A3059) for 1 hour at room temperature. Slides were then incubated with antibodies diluted in 1% goat serum in TBS-T overnight at 4 °C. The following antibodies were used: anti-cytokeratin 14 conjugated to Alexa Flour (AF) 488 (clone LL002, 1:300 dilution, Novus Biologicals, Cat. No. NBP2-34675AF488), anti-cytokeratin 19 (clone EP1580Y, 1:200 dilution, abcam, Cat. No. ab52625), anti-cytokeratin 19 (clone Ba16, 1:100 dilution, abcam, Cat. No. ab20210), anti-CD74 (clone VIC-Y1, 1:50 dilution, Invitrogen, Cat. No. 14-0747-82), and anti-GREB1 (clone PA5-55123, dilution 1:200, Invitrogen, Cat. No. PA5-55123). Slides were washed three times with TBS-T, then incubated with secondary antibodies diluted in 1% goat serum in TBS-T for 30 min at room temperature. The following secondary antibodies were used at a 1:500 dilution: AF647 goat anti-rabbit (Invitrogen, Cat. No. A21244) and AF555 goat anti-mouse IgG1 (Invitrogen, Cat. No. A21127). Slides were washed thrice with TBS-T and dehydrated using a reverse ethanol series. Finally, slides were incubated in xylene for 3 min and then mounted with mounting medium containing DAPI (Invitrogen, Cat. No. P36931). Sections were imaged using an ECHO RVL2-K2 fluorescence microscope. Cells exhibiting nuclear GREB1 staining within luminal epithelial cells were counted independently by two blinded researchers. A minimum of three epithelial regions was counted per sample. To account for differences in staining batches and researcher bias, the percentage of GREB1^+^ cells per sample was divided by total GREB1^+^ cells.

### Flow Cytometry

PDOs were digested to single cells by incubation with TrypLE for 10 min at 37 °C. Single cells were washed with PBS and blocking was performed with 10% goat serum in PBS for 15 min at 4 °C. Cells were then incubated with conjugated antibodies for 45 min at 4 °C. The following antibodies were used at a 1:50 dilution: anti-CD74 conjugated to PE (clone LN2, BioLegend, Cat. No. 326808), anti-CD326/EpCAM conjugated to brilliant violet 421 (clone 9C4, BioLegend, Cat. No. 324220), and anti-CD49f conjugated to AF647 (clone GoH3, BioLegend, Cat. No. 313609). Unstained cells were used as negative control. Cells were washed and filtered through a 40 µm cell strainer (Falcon, Cat. No. 352235). Flow cytometric analysis was performed on an Attune Nxt flow cytometer (Fisher Scientific), and data were analyzed using FlowJo (v10). Cells were first gated based on forward and side scatter (FSC/SSC) to exclude debris. Single cells were then identified by gating on SSC-A versus SSC-H. LASP cells were gated based on co-expression of EpCAM and CD49f.^74,113–116^ The proportion of CD74^+^ cells within the LASP population was gated relative to each sample’s negative control.

### Spatial transcriptomics

Spatial transcriptomics was performed on FFPE sections of six normal human breast tissues using the STOmics Stereo-seq OMNI kit (STOmics, Cat. No. 211SN114 and 111KL160) according to the manufacturer’s instructions. Adjacent FFPE sections were stained with hematoxylin and eosin (H&E) using standard protocols. One sample was excluded due to insufficient library concentrations. Libraries were sequenced on a Complete Genomics DNBseq T7 platform using the 200-cycle configuration at the UCLA Technology Center for Genomics & Bioinformatics. Raw FASTQ files were aligned to the GRCh38 human reference genome, transcripts were quantified, and cells were segmented to generate spatial feature expression matrices using the Stereo-seq Analysis Workflow (SAW) software suite (v8.1.3).^117^ Cells expressing fewer than 3 genes, with fewer than 200 total counts, >15% mitochondrial content, and >500 genes detected were filtered using StereoPy (v1.6.0) and scanpy (v1.11.2).^118,119^ A second sample was filtered out during quality control since it did not pass the filtering criteria. Data from all samples were then merged, and the cell2location pipeline (v0.1.4) was applied using the single-nucleus RNA sequencing dataset as a reference.^120^ Based on cell2location-derived cell type abundance predictions, cells were clustered using StereoSiTE’s cellular neighborhood estimation framework (v2.2.3).^121^ Because cluster 0 was characterized by low transcript levels rather than a distinct gene expression signature, it was re-clustered, resulting in a total of 16 clusters. Figures were generated using matplotlib (v3.10.3) and plotly (v6.5.0). Spatial proximity was assessed by constructing a BallTree (sklearn, v 1.7.0)^122^ on spatial coordinates of immune or epithelial cells and querying neighboring cells within 50 µm. Although HS and LASP cells could be initially identified, their transcriptional profiles showed substantial overlap, consistent with the known similarity between luminal epithelial subtypes. Due to the limited number of genes detected per cells (∼150 genes per cell), these populations were therefore combined into a unified luminal group for downstream analysis. To analyze differences in luminal cells in spatial relation to immune cells, the raw count matrices were exported for analysis in R with Seurat. Data was normalized, variable features were identified, and data was scaled following the default Seurat workflow prior to identifying DEGs between luminal cells distant and proximal to immune cells as described above. For pseudo-bulk analysis, raw gene counts were extracted and summed across cells within each patient to generate pseudo-bulk expression profiles. Differential expression between luminal cells near and far from immune cells of pseudo-bulked samples was performed in R using edgeR (v4.6.3).^123^ Genes with missing values were set to zero, and libraries were normalized using the trimmed mean of M-values (TMM) method. A generalized linear model was fitted, including patient as a blocking factor and proximity group as the variable of interest. Differential expression was assessed using quasi-likelihood F-tests.

### Statistical analysis

Statistical tests that were used to analyze data are noted in the figure legends and methods. In addition, the Shapiro-Wilk test was performed to assess normality of data where appropriate. Correction for multiple testing was performed where appropriate using the Benjamini-Hochberg or Bonferroni method as noted in the figure legends. P values < 0.05 were considered significant. Bar plots show mean ± standard error of the mean (SEM). The number of biological replicates is indicated in the figure and/or figure legend. All experiments were performed in at least technical triplicate.

## APPENDIX

### Lead contact

Requests for further information and resources should be directed to the lead contact, Jennifer Rosenbluth.

## Acknowledgements

We thank all members of the Rosenbluth lab for technical support and the Breast Care Center (BCC) interns for assistance with tissue collection. We are grateful to the UCSF Genomics CoLab, the Chan Zuckerberg Biohub Genomics platform, and the UCLA Technology Center for Genomics & Bioinformatics for assistance with sequencing experiments. Schematic figures were generated using BioRender.com. Funding support was provided by the Doris Duke Charitable Foundation, the V Foundation, the UC STOmics grant program, and the National Institutes of Health R01 CA281361. We thank all patients who agreed to donate tissue for this study.

