## Supplementary Figures for "Hormone signaling and immune programs define differential endocrine responsiveness in high-risk breast tissue"

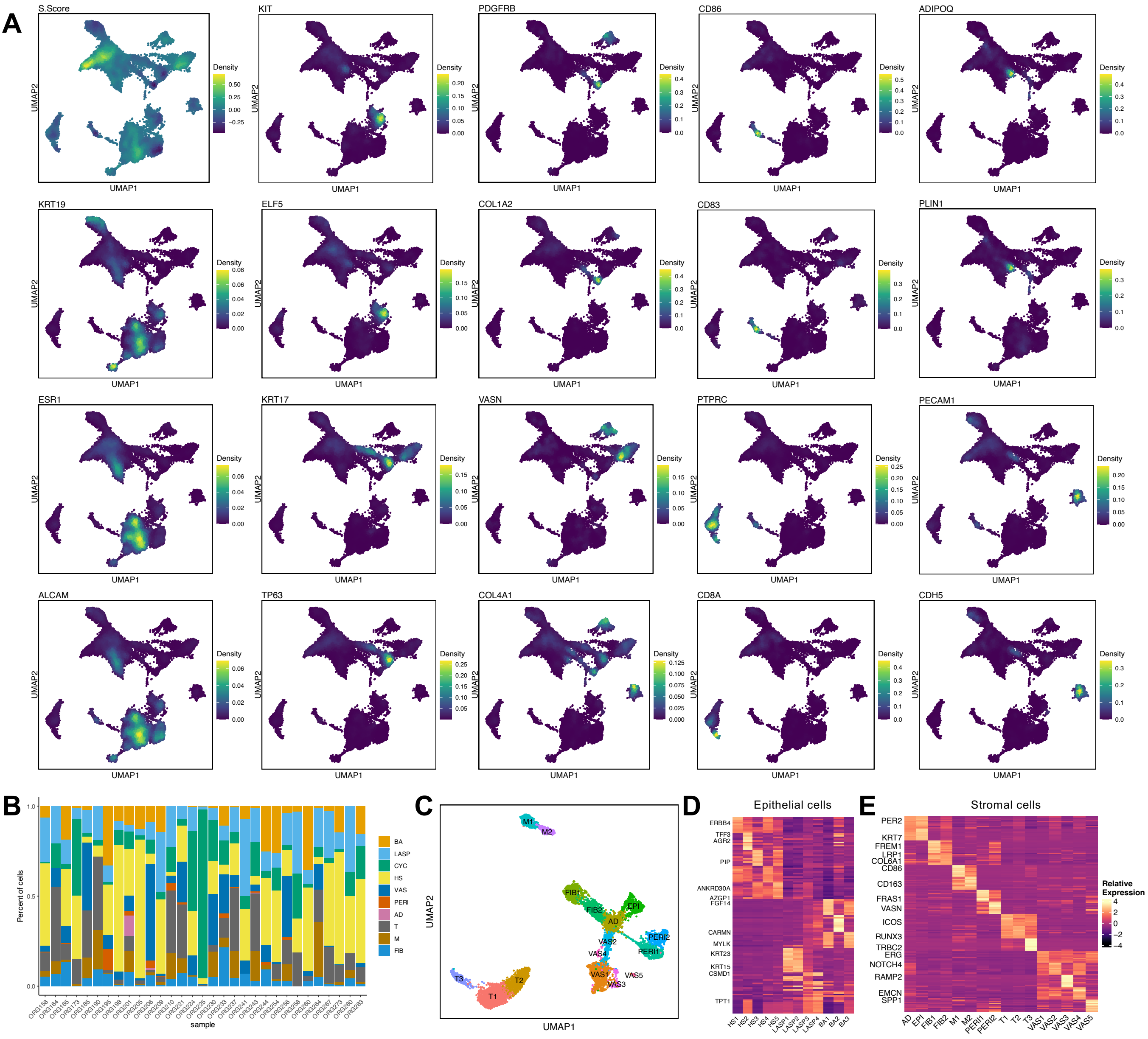


**Supplementary Figure 1: Generation of a snRNA-seq dataset annotated for imaging-based breast cancer risk factors.** A) Expression of lineage-specific cell markers in normal human breast tissues. Density plots showing expression of representative cell type-specific genes for cycling cells (S score), luminal epithelial cells (*KRT19*), HS cells (*ESR1*, *ALCAM*), LASP cells (*KIT*, *ELF5*), basal cells (*KRT17*, *TP63*), fibroblasts (*PDGFRB*, *COL1A2*), pericytes (*VASN*, *COL4A1*), macrophages (*CD86*, *CD83*), T cells (*PTPRC*, *CD8A*), adipocytes (*ADIPOQ*, *PLIN1*), and vascular cells (*PECAM1*, *CDH5*) . B) Distribution of cell types across individual samples. Stacked bar plot illustrating cellular composition (color-coded) per sample. C) Clustering of cell populations of the normal breast stroma. UMAP plot showing 7,457 stromal cells after re-clustering reflecting one or more cell states of adipocytes, fibroblasts, immune cells (macrophages and T cells), pericytes, and vascular cells. D) Expression of cluster-specific differentially expressed genes in epithelial cells of the normal human breast. Heatmap showing expression of the top ten highest expressed genes per cluster in epithelial cells. E) Expression of cluster-specific differentially expressed genes in stromal cells of the normal human breast. Heatmap showing expression of the top ten highest expressed genes per cluster in stromal cells.

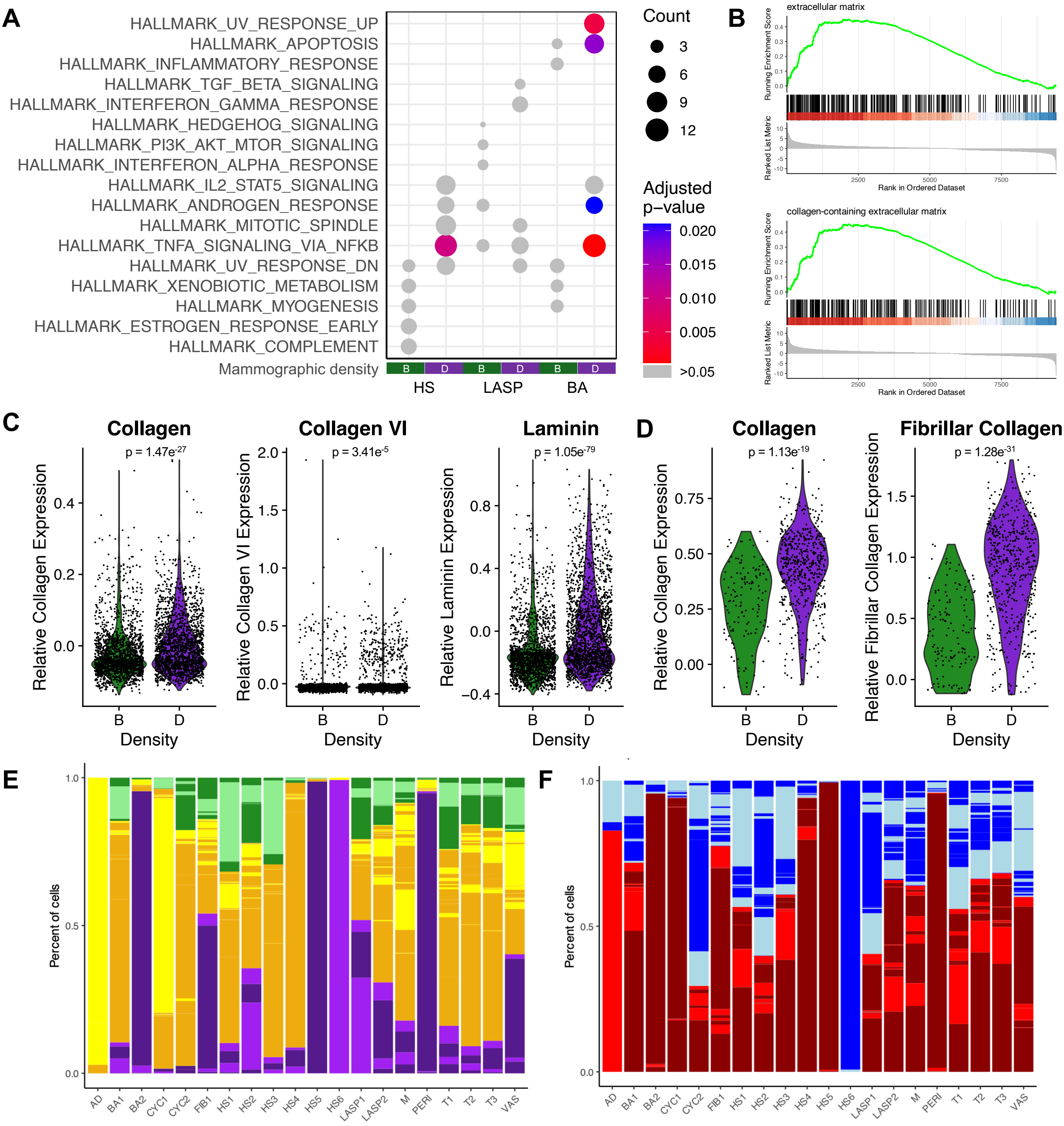


**Supplementary Figure 2: Extracellular matrix and collagen gene expression are increased in extremely dense breast tissue.** A) Differently enriched pathways in epithelial cells from tissues with distinct mammographic densities. Dot plot showing pathways in epithelial cells from tissues with distinct mammographic densities. B) Enrichment of gene sets associated with the extracellular matrix and collagen in dense breast tissues. Gene set enrichment plots showing enrichment of extracellular matrix (top) and collagen-containing extracellular matrix (bottom) gene sets in DEGs of tissues with extreme versus scattered MD. C) Expression of extracellular matrix genes in epithelial cells of the normal human breast. Violin plots showing relative expression scores of genes encoding collagens, collagen IV family members, and laminins, in tissues with scattered (B, green) or extreme (D, purple) MD. P values by Wilcoxon rank sum test with Benjamini Hochberg correction. D) Expression of extracellular matrix genes in fibroblasts of the normal human breast. Violin plots showing relative expression scores of genes encoding collagens and fibrillar collagens in fibroblasts from tissue with scattered (B) and extreme (D) MD. P values by Wilcoxon rank sum test with Benjamini Hochberg correction. E) Distribution of samples with distinct MDs across cell types. Stacked bar plot showing proportion of samples with distinct MDs (green = B, orange = C, purple = D) per cell type. Different shades demark different samples. F) Distribution of samples with distinct BPE levels across cell types. Stacked bar plot showing proportion of samples with distinct BPE levels (red = high, blue = low) per cell type. Different shades demark different samples.

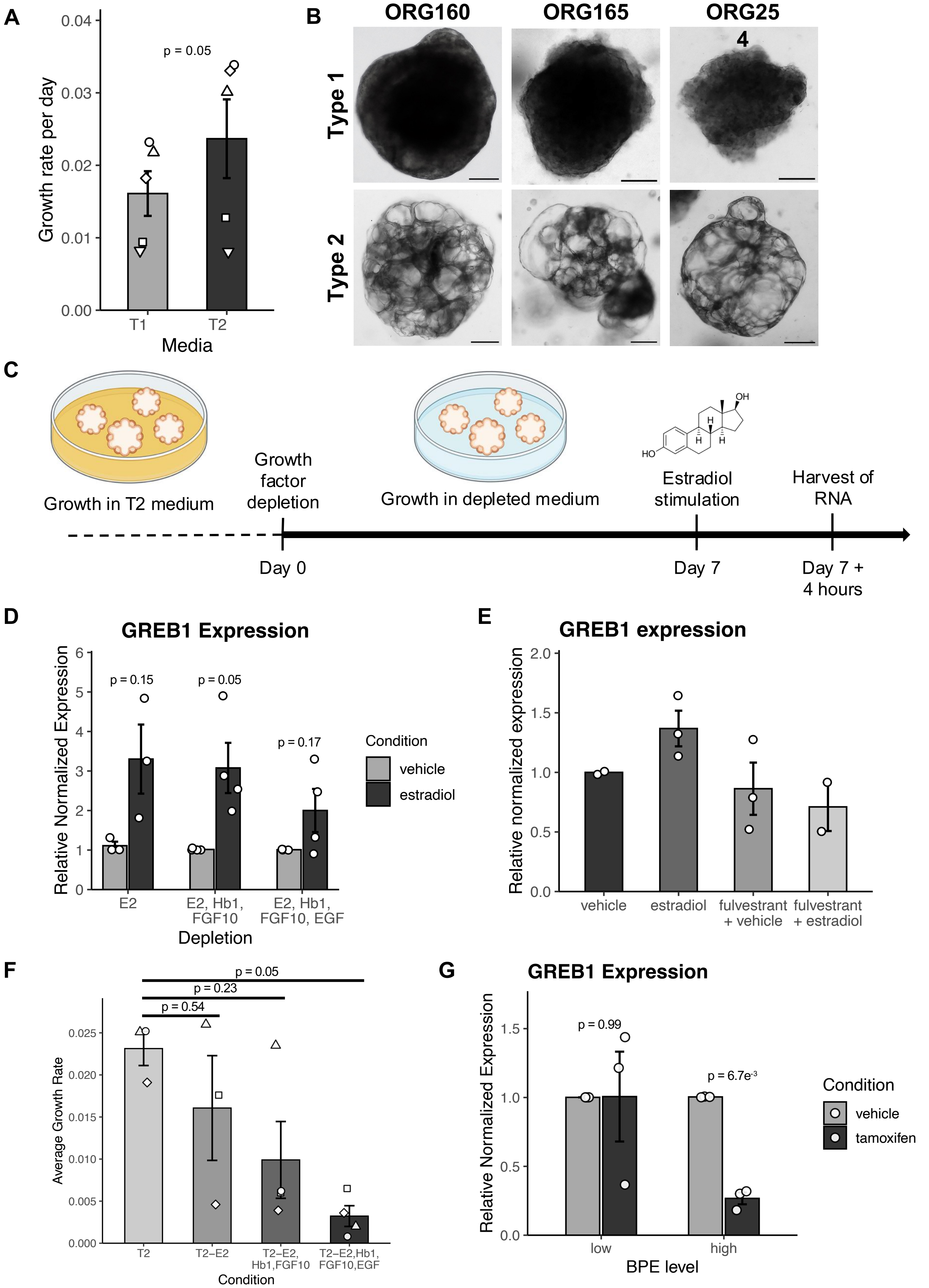


**Supplementary Figure 3. PDO lines from tissues with distinct BPE levels reflect changed estrogen responsiveness.** A) Growth rates of PDOs in different media types. Bar plot showing growth rates of PDOs grown in type 1 or type 2 medium. Shapes indicate different samples. P value by paired t test, n = 5. B) PDOs morphology in different media types. Representative bright field images showing three distinct PDOs grown in type 1 (top) or type 2 (bottom) organoid medium. Scale bar = 100 µm. C) Schematic overview of the protocol used to assess PDO responsiveness to β-estradiol stimulation. D) Expression of *GREB1* in response to β-estradiol after depletion of different growth factor combinations. Bar plot showing expression of estrogen-response gene *GREB1* in organoids that were depleted of β-estradiol (E2, n = 3), and/or heregulin β1, fibroblast growth factor 10 (FGF10) (n = 4), and/or epidermal growth factor (EGF), n = 4) for seven days before stimulation with 100 nM β-estradiol for four hours. P values by paired t test with Benjamini Hochberg correction. E) Expression of *GREB1* in PDOs after treatment with the SERD fulvestrant. Bar plot showing expression of estrogen response gene *GREB1* in a PDO after stimulation with vehicle control, 100 nM β-estradiol, 10 nM fulvestrant and vehicle control, or 10nM fulvestrant and 100 nM β-estradiol. No significant differences were detected by t-test, n = 1 PDO in triplicate. F) Growth rates of PDOs in response to depletion of different growth factor combinations. Bar plot showing average growth rate of PDOs grown in full medium (no depletion, T2) and medium that was depleted of β-estradiol (E2), and/or heregulin β1, fibroblast growth factor 10 (FGF10), and/or epidermal growth factor (EGF). P values by paired t test with Benjamini Hochberg correction. G) Expression of *GREB1* in PDOs after treatment with tamoxifen. Bar plot showing expression of estrogen-response gene *GREB1* in PDOs from tissues with distinct BPE levels treated with vehicle control or 1 µM tamoxifen. P value by t test with Benjamini Hochberg correction, n = 3.

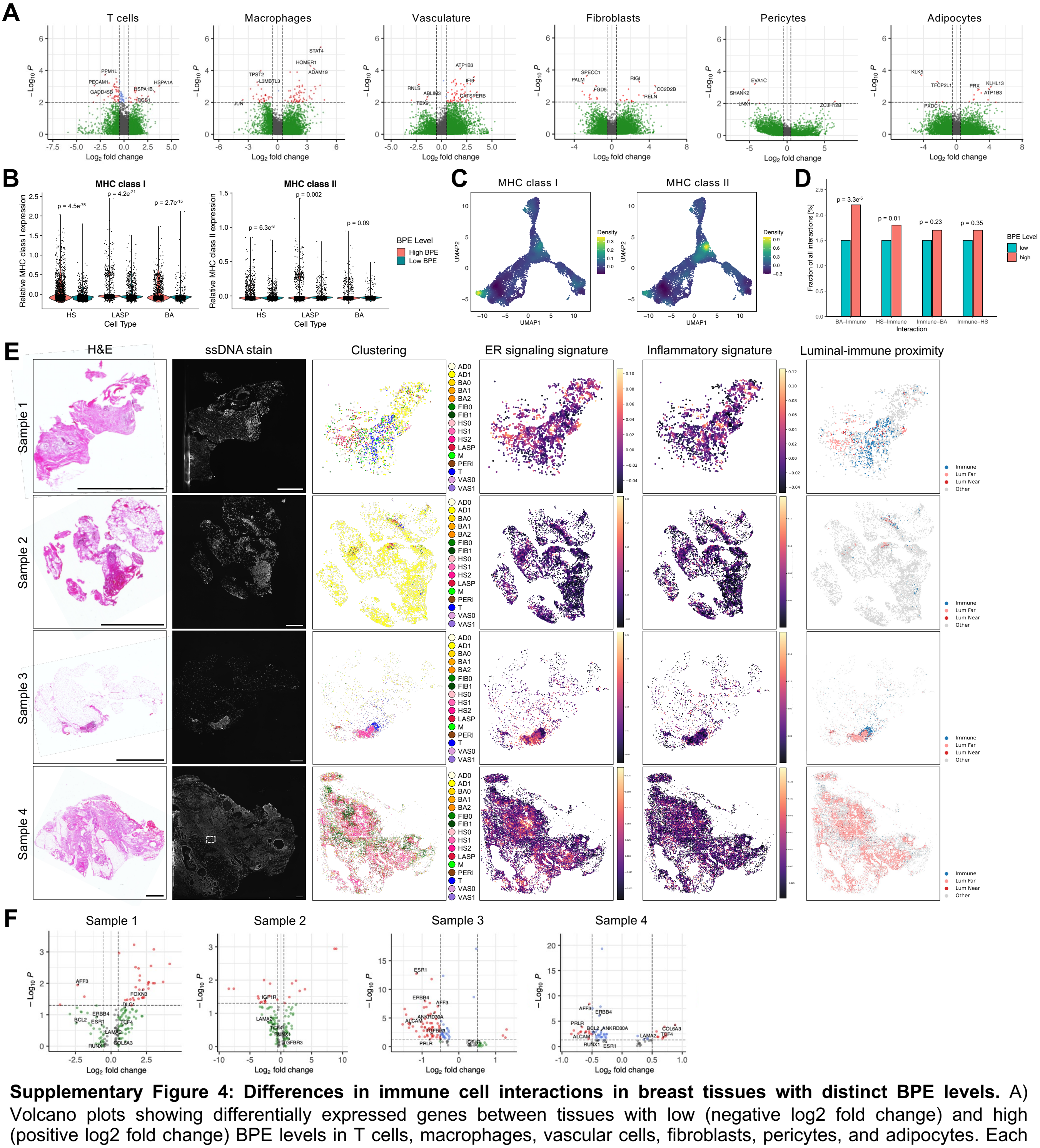


**Supplementary Figure 4: Differences in immune cell interactions in breast tissues with distinct BPE levels.** A) Volcano plots showing differentially expressed genes between tissues with low (negative log2 fold change) and high (positive log2 fold change) BPE levels in T cells, macrophages, vascular cells, fibroblasts, pericytes, and adipocytes. Each dot represents a gene and selected genes are labelled. Red = p value < 0.01 and log2 fold change > 0.5; blue = p value < 0.01 and log2 fold change < 0.5; green = p value > 0.01 and log2 fold change > 0.5; grey = p value > 0.01 and log2 fold change < 0.5. B) Expression levels of MHC genes in epithelial cells of the normal human breast. Violin plot showing scores of antigen-presentation gene signatures in the three main epithelial populations in tissues with distinct BPE levels. P values by Wilcoxon rank sum test with Benjamini Hochberg correction. C) Expression of MHC genes in epithelial cells of the normal human breast. Density plots showing expression of genes encoding MHC class I (left) and MHC class II (right) proteins in epithelial cells of the normal human breast. D) Interactions between HS and basal with immune cells in tissues with distinct BPE levels. Bar plot showing percentage of total cell-cell interactions detected that account for interactions between the basal (BA) and hormone-sensing (HS) cells and immune cells (macrophages and T cells) in tissues with distinct BPE levels. P values by 2-sample test for equality of proportions with continuity correction. E) Spatial transcriptomics images from four normal human breast samples showing hematoxylin and eosin staining (H&E, scale bar = 2000 μm), single-stranded DNA (ssDNA) staining (scale bar = 500 μm), spatial clustering, estrogen receptor (ER) signaling signatures, inflammatory signatures, and proximity (50 μm) of luminal epithelial cells (red) to immune cells (blue). White insert in the ssDNA panel, sample 4, indicates zoom-in shown in Figure 4. F) Gene expression differences between luminal epithelial cells close and far from immune cells in spatial transcriptomics data. Volcano plot showing adjusted p values and log2FC of DEGs between luminal cells within 50 μm (positive log2FC) versus distant (negative log2FC) from immune cells in for each sample of the spatial transcriptomics data.

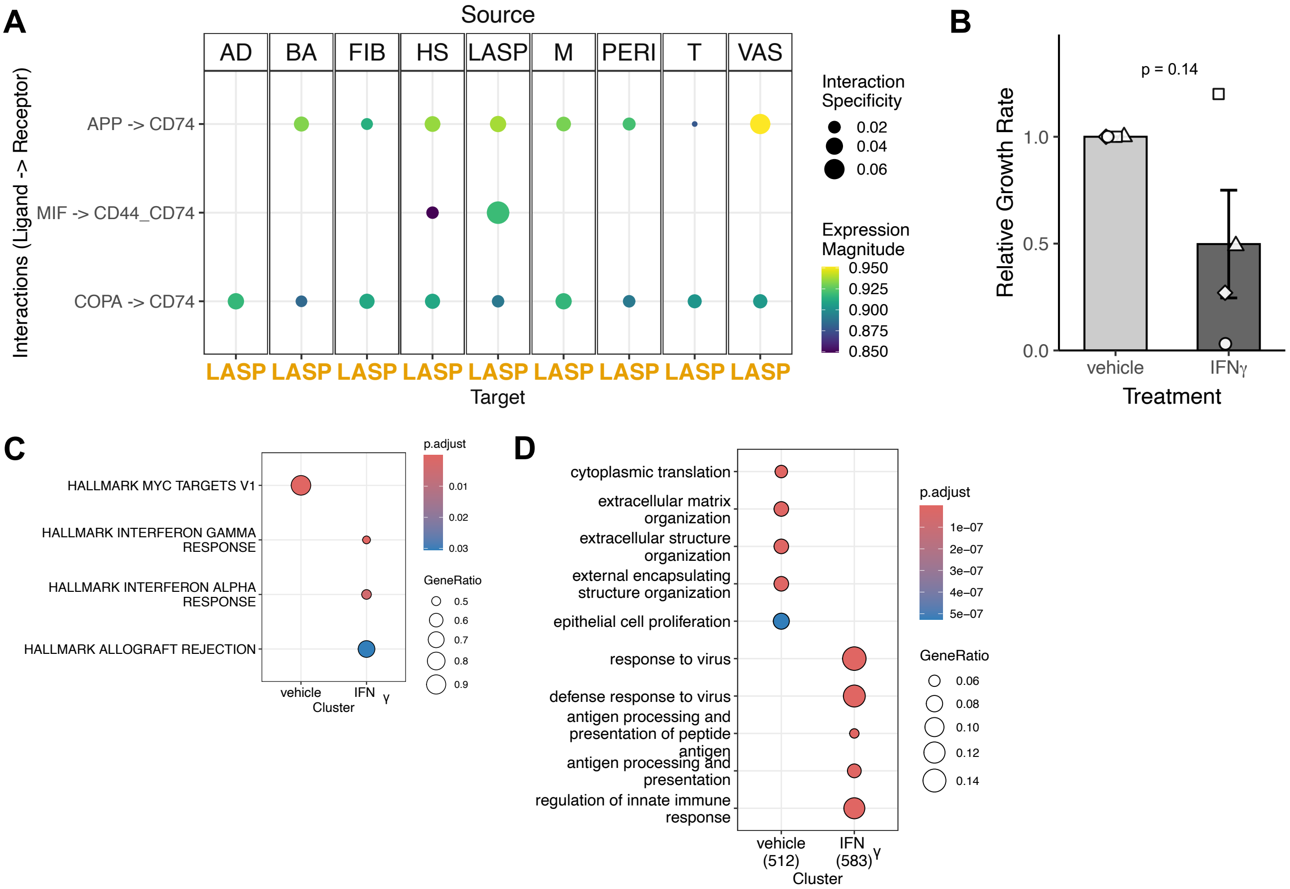


**Supplementary Figure 5. IFNγ treatment induces gene expression changes in normal breast PDOs.** A) Expression of CD74 and its ligands in LASP cells of tissues with high BPE levels. Dot plot showing interactions involving CD74 expressed on LASP cells with all other cell types. B) Growth rate of PDOs treated with IFNγ. Bar plot showing relative growth rate of PDOs treated with vehicle control or 5 ng/ml IFNγ for seven days. P value by paired t test. C) Enriched hallmark gene signatures in PDOs treated with IFNγ. Dot plot showing hallmark gene sets enriched in PDOs treated with vehicle control or 5 ng/ml IFNγ. D) Enriched gene ontology gene signatures in PDOs treated with IFNγ. Dot plot showing Gene Ontology Cellular Component (GO-CC) gene sets enriched in PDOs treated with vehicle control or 5 ng/ml IFNγ.
